# Gene co-expression analysis of human *RNASEH2A* reveals functional networks associated with DNA replication, DNA damage response, and cell cycle regulation

**DOI:** 10.1101/2020.08.27.270595

**Authors:** Stefania Marsili, Ailone Tichon, Francesca Storici

**Affiliations:** School of Biological Sciences, Georgia Institute of Technology, Atlanta, GA 30332

## Abstract

Ribonuclease H2 (RNase H2) is a key enzyme for the removal of RNA found in DNA-RNA hybrids, playing a fundamental role in biological processes such as DNA replication, telomere maintenance and DNA damage repair. RNase H2 is a trimer composed of three subunits, being RNASEH2A the catalytic subunit. *RNASEH2A* expression levels have been shown to be upregulated in transformed and cancer cells. In this study we used a bioinformatics approach to identify *RNASEH2A* co-expressed genes in different human tissues to uncover biological processes in which *RNASEH2A* is involved. By implementing this approach, we identified functional networks for *RNASEH2A* that are not only involved in the processes of DNA replication and DNA damage response, but also in cell cycle regulation. Additional examination of protein-protein networks for RNASEH2A by the STRING database analysis, revealed a high co-expression correlation between *RNASEH2A* and the genes of the protein networks identified. Mass spectrometry analysis of RNASEH2A-bound proteins highlighted players functioning in cell cycle regulation. Further bioinformatics investigation showed increased gene expression of *RNASEH2A* in different types of actively cycling cells and tissues, and particularly in several cancers, supporting a biological role for RNASEH2A, but not the other two subunits of RNase H2, in cell proliferation.

## Introduction

The development of high-throughput tools to monitor gene expression levels in a specific cell type and tissue has allowed the characterization of gene expression patterns throughout the human body. Such approach was utilized in several studies to identify molecular signatures of tissues and cells [1,2,3,]. Genes that share a molecular pathway in a given tissue or cell should be co-expressed in a spatial and temporal manner. Therefore, examining genes that show high co-expression correlation in multiple tissues can be used to identify a shared molecular pathway between those genes. The Genotype-Tissue Expression portal (GTEx) [3] is a suitable platform to identify gene co-expression in the human body based on the availability of the full transcriptome in 53 different human tissues. Gene ontology analysis of the co-expressed genes might shed light on genes with uncharacterized functions and help discover additional functions of genes for which their role is partially known. RNASEH2A, together with RNASEH2B and RNASEH2C, composes the holoenzyme RNase H2 [4,5]. RNASEH2B and RNSEH2C are used as the scaffold for RNASEH2A, which serves as the catalytic subunit of the RNase H2 complex [6]. The function of RNase H2, together with RNASEH1, is to cleave RNA of RNA-DNA hybrids, which can be formed during transcription [7], DNA replication [8] and repair [9]. RNA-DNA hybrids, if not repaired, can harm the genomic DNA by different mechanisms, such as modifying the DNA structure, blocking DNA replication and transcription, causing hyperrecombination and mutation or chromosome loss [10,11,12, 13, 14].

Phenotypically, it has been postulated that the RNA-DNA hybrids mimic the infection of nucleic acids of viral origin, thus activating the innate immune response [15]. An example of a pathologic condition is the Aicardi-Goutieres Syndrome (AGS), a severe autosomal recessive neurological disorder with symptoms similar to *in-utero* viral infection [16,17]. Genetic studies have linked AGS to mutations in the genes composing RNase H2. However, no evidence was found that RNASEH1 is linked to the AGS pathology [15], highlighting a marked difference between the function of these two enzymes. In immune-precipitation pulldown experiments it was shown that RNASEH2B and RNASEH2C interact with each other and with RNASEH2A, while RNASEH2A was found in an unbound fraction as well [6]. Another distinctive feature of *RNASEH2A* comes from examining its expression levels in cancer compared to normal tissues. A previous study showed that in human mesenchymal stem cells transformed by the over-expression of several oncogenes, *RNASEH2A* was among the genes with the highest and earliest up fold change in their expression [18]. In addition, increased levels of *RNASEH2A* in cancer compared to normal tissues and cells were also reported in cervical cancer [19], prostate cancer [20], colorectal carcinoma [21] and triple negative breast cancer [22]. In support of this finding, a recent study by Luke’s group [23] highlights a differential cell-cycle regulation for the activity and level of the *RNASEH2A* orthologous gene in yeast *Saccharomyces cerevisiae* (*RNH201*), peaking at S and G2 phases of the cell cycle, supporting the hypothesis that RNASEH2A plays a role in cancer progression. This is in contrast with the yeast *RNASEH1* ortholog, *RNH1*, the activity and level of which is not altered throughout the cell cycle [23]. With the goal to better understand the biological role of RNASEH2A, we further characterized the expression level of *RNASEH2A* in different human tissues, hypothesizing that, identifying patterns of its expression, can shed new insights on its function.

## Results

### *RNASEH2A* is co-expressed with genes that function in cell cycle regulation

To study RNASEH2A function based on co-expressed genes we developed an approach (Fig. 1a) that utilizes data obtained from GTEx portal [3] consisting of RNA-seq output of 42,548 nuclear and mitochondrial genes including coding and non-coding and isoforms that are expressed in at least one out of the 53 human tissues (raw data in Supplementary Table S1). Then, we examined the gene ontology of the top 2% co-expressed correlated genes (851 genes) using the GOrilla analysis tool [24]. We observed a significant enrichment of genes involved in the biological processes of cellular response to DNA damage stimulus (p=2.77E-05) and DNA replication (p=2.66E-04), compatible with the known functions of RNASEH2A. In addition, we found enrichment of genes involved in mitotic cell-cycle process (p=2.80E-14), regulation of microtubule cytoskeleton organization (p=1.39E-07), and chromosome segregation (p=9.00E-04) (Table 1 and full data in Supplementary Table S2). For molecular function, we found enrichment of catalytic activity on DNA (p=9.59E-06), but also microtubule binding (p=4.25E-05) and kinase binding (p=3.28E-04) (Table 1 and full data in Supplementary Table S2). For cellular component, we found genes enrichment in chromosomal part (4.06E-08), spindle pole (p=1.13E-06), as well as nucleoplasm (p=3.52E-06), midbody (p=5.01E-06), microtubule organizing center (p=5.13E-06), and condensed chromosomes outer kinetochore (p=8.72E-06) (Table 1 and full data in Supplementary Table S2). Based on these results, we suggest that *RNASEH2A* is involved, among others, in three main functional networks related to DNA replication, DNA damage repair and regulation of chromosome segregation in mitosis. To verify these results, we utilized a different approach by implementing protein network association data obtained from STRING protein-protein association network v11[25] on the *RNASEH2A* co-expression correlation analysis (Figure 1b). We were interested in proteins found to interact and function with *RNASEH2A* and/or with its top co-expressed correlated genes, and in their co-expression analysis correlation with *RNASEH2A*. We identified the top 20 known and characterized protein interactors for each of the top 9 *RNASEH2A* co-expressed genes (including *RNASEH2A* itself), (full list of proteins in Supplementary Table S3). When those proteins were aligned with the co-expressed correlated *RNASEH2A* genes, we found that out of the 139 proteins, 69 occurred among the top 2% co-expressed correlated genes, 83 proteins in the top 5% and 98 proteins in the top 10% (full data in Supplementary Table S4). These results indicate that the analysis of genes co-expression correlation with *RNASEH2A* can also be applied to its protein translated form.

**Figure 1.**
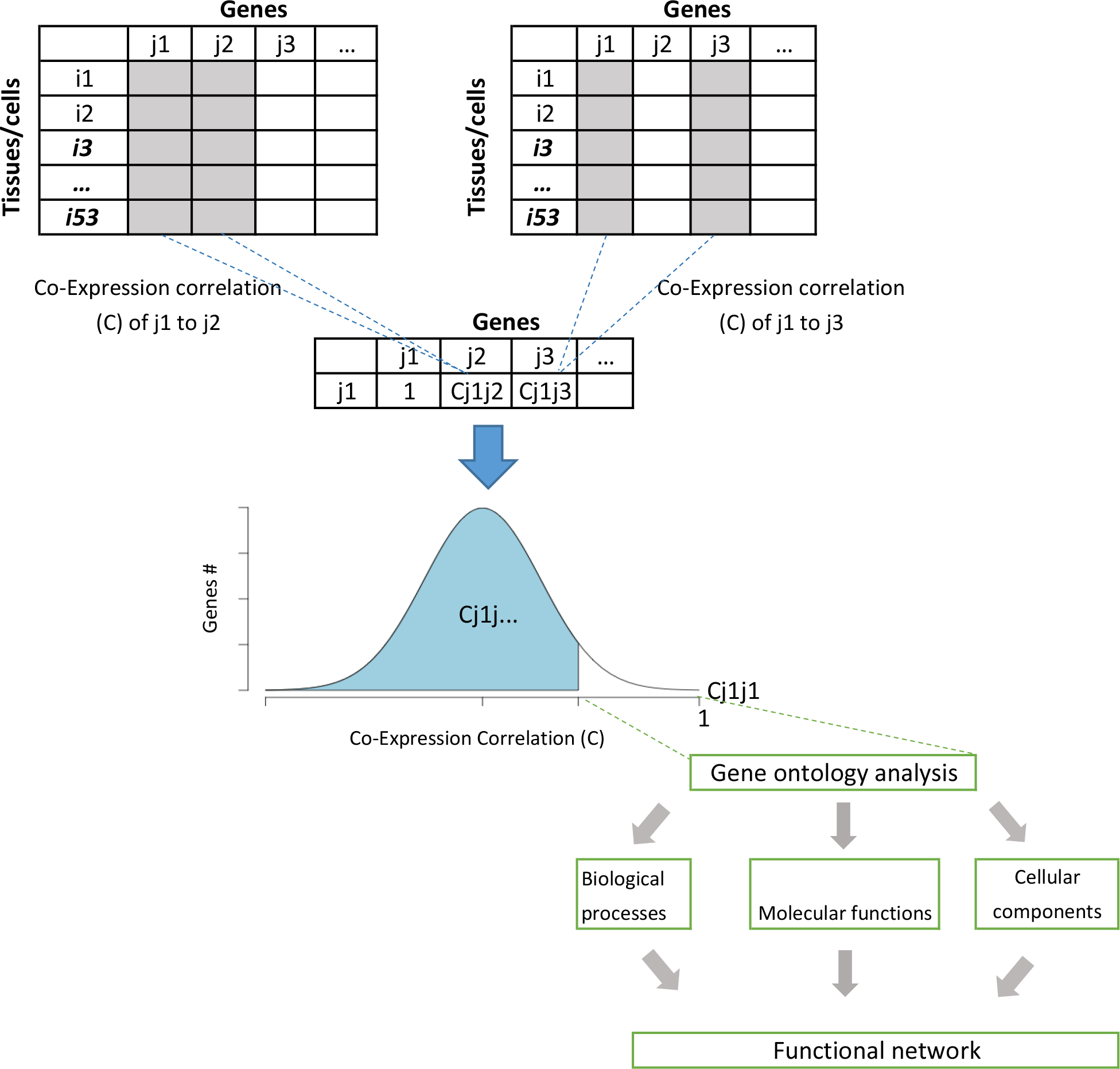
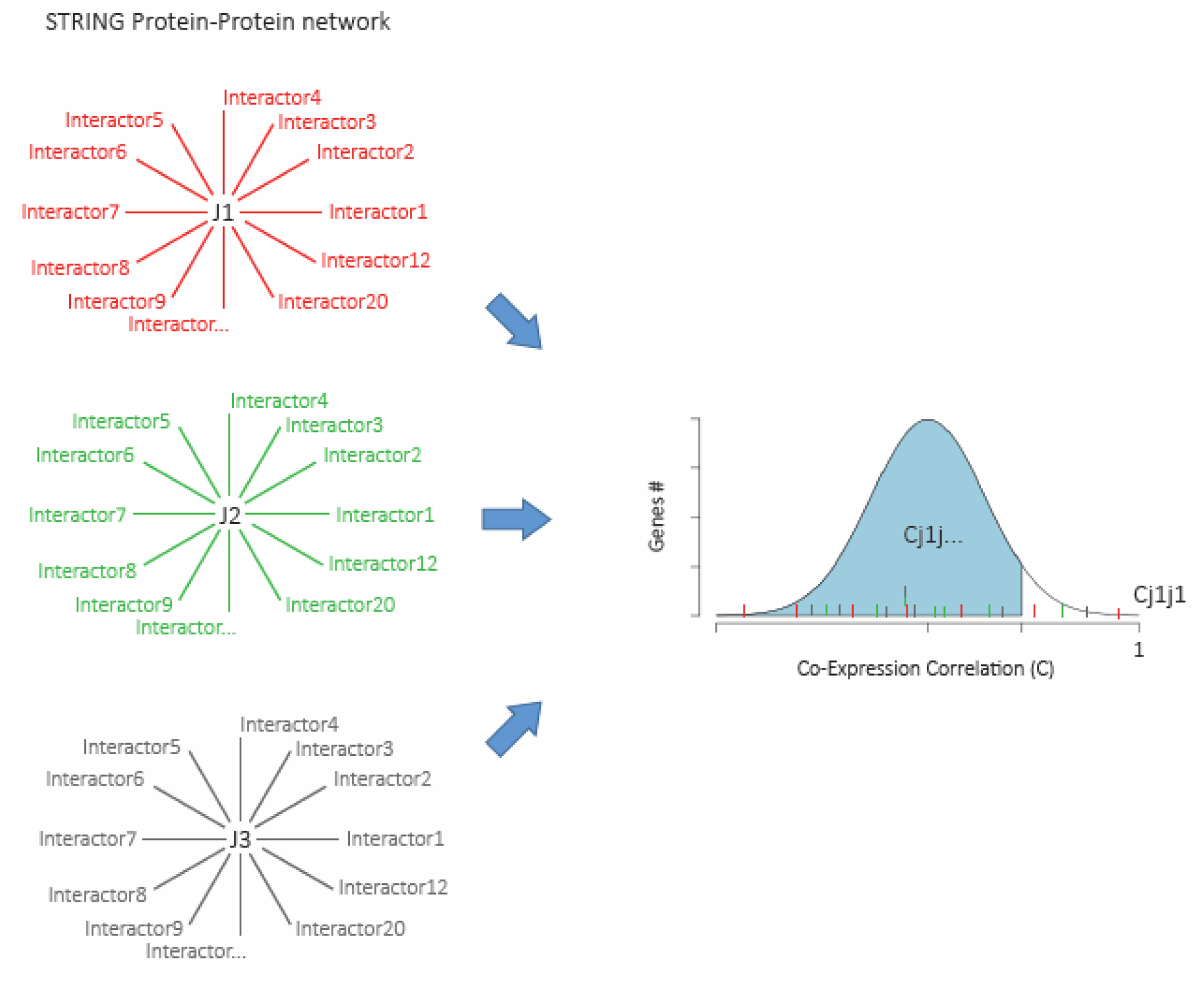
Co-expression correlation analysis approach. **1a.** Diagram showing the process of analyzing gene functional networks using co-expression correlation analysis. The analysis is based on any database that show expression levels on multiple tissues and cells from the same organism. **1b**. Diagram showing the process of verifying functional network of genes using proteins associated network. Example data were obtained from STRING protein: protein interactions. J1-3 are proteins for which binding partners were aligned with the Co-Expression Correlation of their genes to gene J1.

**Table 1.**
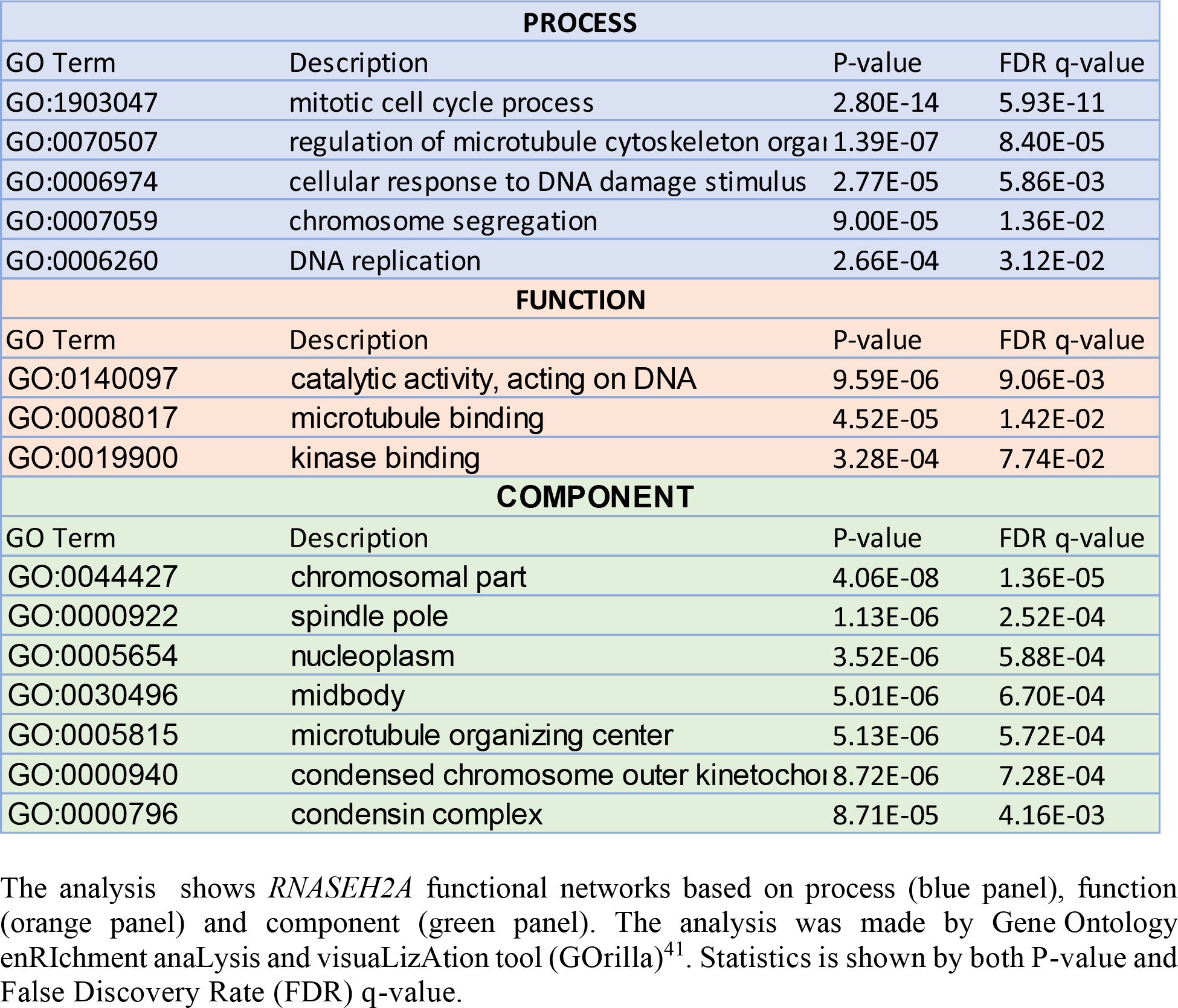
GO term analysis of the top 2% co-expressed genes of *RNASEH2A* reveals possible role of *RNASEH2A* in cell cycle regulation.

### Mass spectrometry analysis supports a functional role of RNASEH2A in mitosis regulation

To identify proteins that interact with RNASEH2A, we over-expressed eGFP-RNASEH2A and the control eGFP plasmids in HEK293 cells. We performed co-immunoprecipitation (Co-IP) using an anti-GFP antibody (Figure 2a). We also confirmed that both GFP and RNASEH2A antibodies identify the same GFP-RNASEH2A protein (Figure 2b). After confirming the successful pulldown of RNASEH2A protein, we analyzed the eluted proteins that were interacting with RNASEH2A *via* mass spectrometry. The analysis revealed a short list of 22 human proteins with at least 3 peptides peptide spectrum matches (PSMs) interacting with RNASEH2A (Table 2 and full data in Supplementary Table S5). RNASEH2B and RNASEH2C were interacting with RNASEH2A with 27 and 17.5 PMSs identified respectively, compared to none observed in the control, confirming the validity of the Co-IP experiment. Interestingly, among the proteins interacting with RNASEH2A we found the T-complex protein 1 subunits theta (CCT8) and beta (CCT2), the Carbamoyl-phosphate synthetase 2-aspartate transcarbamylase-dihydroorotase (CAD), Nucleolin (NCL) and the Heat shock protein Hsp90-beta (HSP90AB1), with a co-expression correlation of 0.840, 0.816, 0.768, 0.791 and 0.505 respectively. All of these RNASEH2A interactors have shown to play a role in cell cycle regulation and mitosis [26,27,28,29].

**Figure 2.**
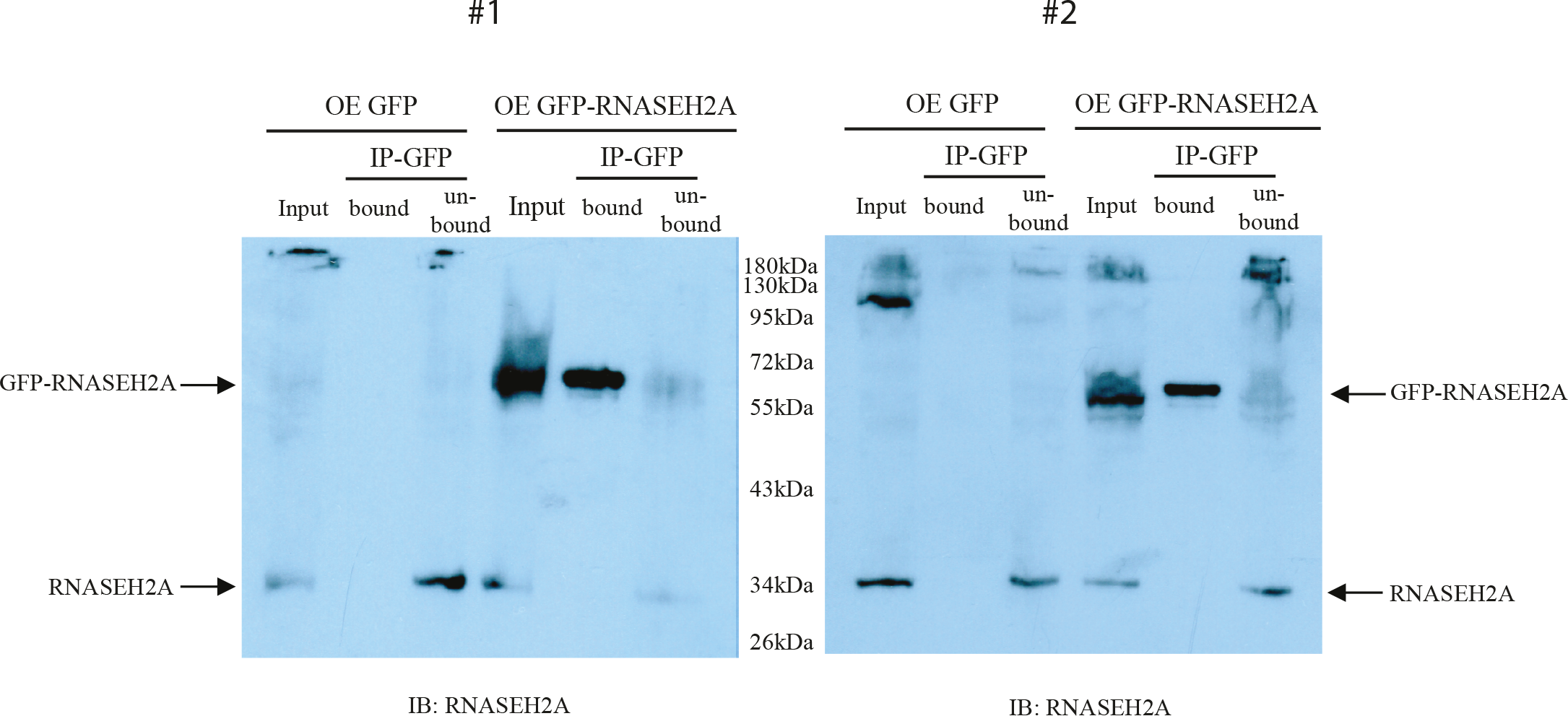
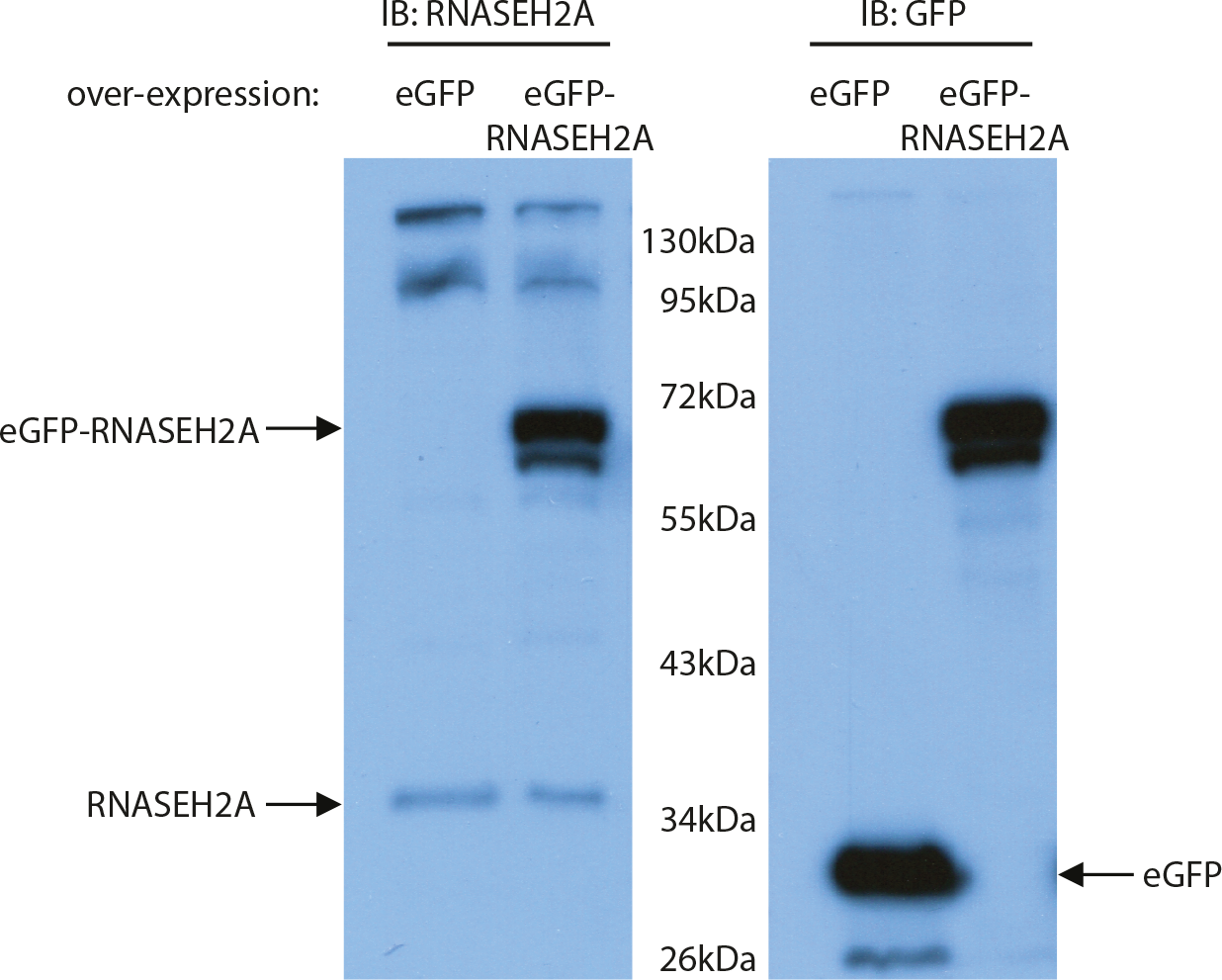
2a. Co-immunoprecipitation (Co-IP) of RNASEH2A. Western Blot of the GFP co-Immunoprecipitation. The two panels indicate two independent experiments. HEK293 cells were OE with the eGFP vector (three lanes from the left in each panel) or with the eGFP-RNAaseH2A vector (three lanes from the right in each panel). IP: anti-GFP. IB: anti-RNASEH2A. Arrows indicate endogenous RNASEH2A (lower arrow) and exogenous eGFP-RNASEH2A (upper arrow). **2b. RNASEH2A Antibody verification.** WB of the anti-RNASEH2A (left panel) and anti-GFP (right panel). In the two panels HEK293 cells were OE with eGFP vector (left lane) or with eGFP-RNASEH2A (right lane). In the left panel arrows indicate endogenous RNASEH2A (lower arrow) and exogenous eGFP-RNASEH2A (upper arrow). In the right panel arrows indicate eGFP (lower arrow) and eGFP-RNASEH2A (upper arrow).

**Table 2.**
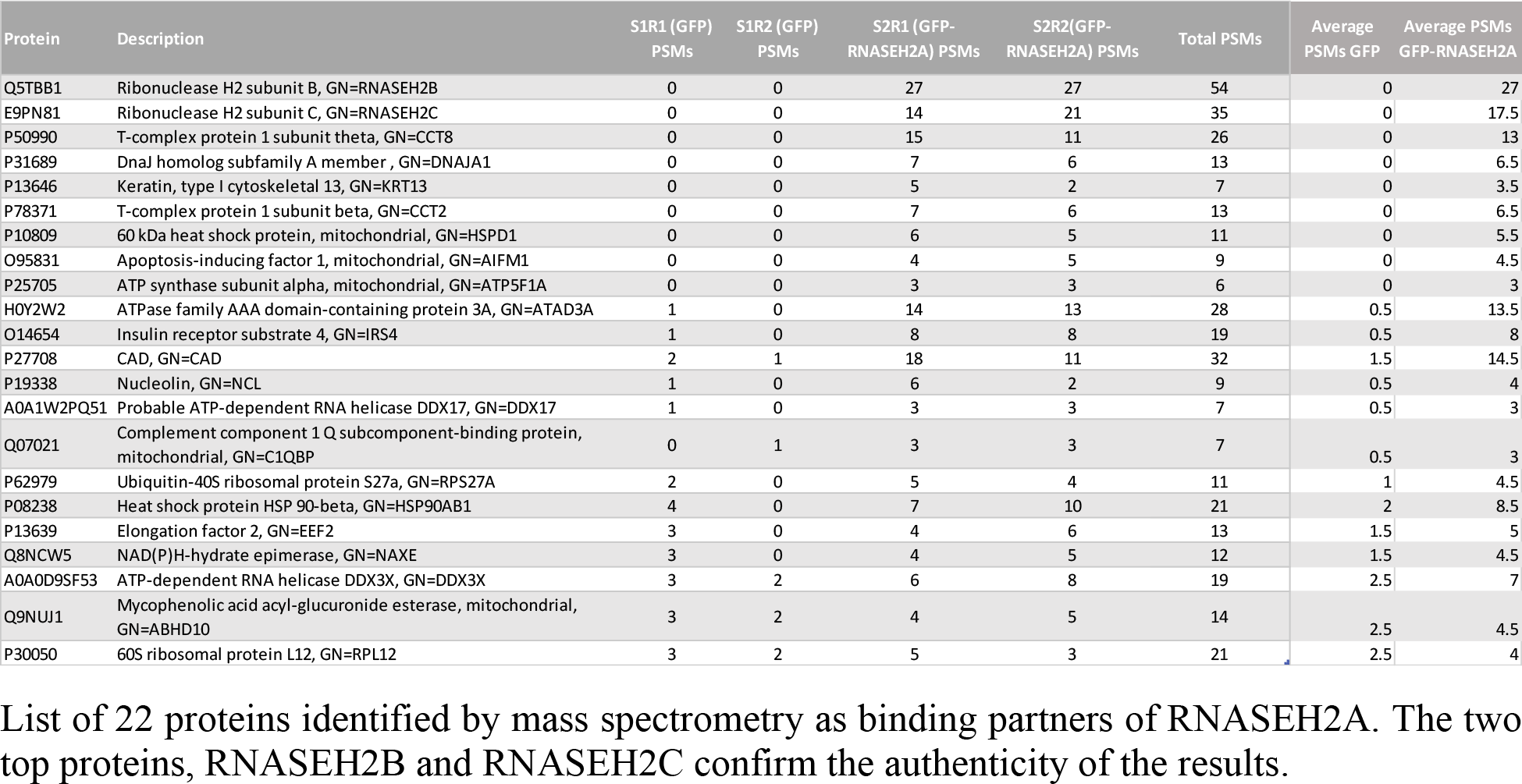
Mass spectrometry analysis reveals interactors of RNASEH2A with role in cell cycle regulation and mitosis.

### *RNASEH2A* expression is increased in actively cycling cells and tissues

Next, we studied the expression levels of *RNASEH* genes using the GTEx v.7 database, which allowed us to compare the expression levels of *RNASEH1*, *RNASEH2A*, *RNASEH2B* and *RNASEH2C* genes in 53 different human tissues including 2 cell lines (Table 3). We noticed that tissues that are part of the reproduction system (such as testis, uterus and cervix) as well as transformed lymphocytes and transformed fibroblasts tend to have a higher expression levels of *RNASEH* genes, while low levels were found mainly in tissues with low proliferation and cell turnover capacity, such as most regions of the brain, kidney, heart and whole blood (Table 3). We then examined the expression level of the *RNASEH* genes in cancer using data obtained from Cancer RNA-seq Nexus (CRN) [30] in 29 different randomly selected tumors in different tissues at different stages of cancer progression compared to their healthy controls (complete list of tumors in Supplementary Table S6 and raw data in Supplementary Table S7). To validate our approach, we also examined the expression level of *MYBL2* and *SCARA5* genes, which are reported to be upregulated and downregulated in cancer, respectively [31]. As expected, *MYBL2* was mostly upregulated in cancer such as triple negative breast cancer, bladder urothelial carcinoma stage 2,4 and lung adenocarcinoma stage 2,4; while *SCARA5* was downregulated in cancer such as bladder urothelial carcinoma stage 2,4; thyroid carcinoma stage 1 and rectum adenocarcinoma stage 2A (Figure 3). *RNASEH2A* was the only *RNASEH* gene that showed increased expression levels in all 29 tumors relative to the control tissues (Figure 3). To study the levels of *RNASEH* genes throughout the progression of cancer, we examined the expression levels of the genes at different stages of lung squamous adenocarcinoma, breast invasive carcinoma and bladder urothelial carcinoma. In all these cancers *RNASEH2A* was upregulated at early stage and remained high with the progression of the disease (Figure 4 and Supplementary Table S8 and raw data in Supplementary Table S9). No other gene, beside *MYBL2*, was upregulated at the different stages studied. These results imply that *RNASEH2A* might have a role in cell proliferation that is dependent on its expression level.

**Table 3.** *RNASEH2A* is highly expressed in human proliferative tissues.

|  | *RNASEH1* | *RNASEH2A* | *RNASEH2B* | *RNASEH2C* |  | *RNASEH1* | *RNASEH2A* | *RNASEH2B* | *RNASEH2C* |
| --- | --- | --- | --- | --- | --- | --- | --- | --- | --- |
| *Liver* | 5.193 | 3.947 | 3.878 | 11.57 | *Adipose - Subcutaneous* | 13.845 | 15.96 | 16.585 | 30.145 |
| *Muscle - Skeletal* | 11.72 | 5.9535 | 5.51 | 9.215 | *Lung* | 12.92 | 14.51 | 19.34 | 30.99 |
| *Heart - Left Ventricle* | 7.251 | 4.262 | 8.466 | 12.71 | *Breast - Mammary Tissue* | 12.93 | 16.39 | 16.65 | 33.895 |
| *Pancreas* | 5.3175 | 6.44 | 10.175 | 11.48 | *Pituitary* | 8.271 | 11.02 | 14.5 | 52.95 |
| *Whole Blood* | 2.726 | 7.958 | 6.471 | 16.33 | *Vagina* | 12.08 | 20.33 | 16.85 | 39.11 |
| *Brain - Putamen (basal ganglia)* | 4.6985 | 5.5965 | 7.875 | 15.635 | *Esophagus - Muscularis* | 14.91 | 8.687 | 22.84 | 46.885 |
| *Brain - Caudate (basal ganglia)* | 5.1025 | 6.1935 | 9.2325 | 17.175 | *Artery - Tibial* | 16.91 | 8.638 | 17.8 | 50.75 |
| *Brain - Substantia nigra* | 6.0725 | 7.4295 | 8.123 | 16.355 | *Thyroid* | 12.615 | 16.935 | 24.905 | 43.205 |
| *Brain - Amygdala* | 5.0275 | 7.4755 | 8.2985 | 17.285 | *Esophagus - Gastroesophageal Junction* | 13.75 | 8.3635 | 22.05 | 53.92 |
| *Brain - Hippocampus* | 5.907 | 6.655 | 7.567 | 18.76 | *Bladder* | 15.4 | 15.95 | 23.68 | 43.32 |
| *Brain - Anterior cingulate cortex (BA24)* | 5.921 | 6.326 | 9.554 | 19.13 | *Artery - Coronary* | 13.65 | 10.5 | 18.58 | 56.4 |
| *Brain - Hypothalamus* | 6.903 | 7.451 | 8.639 | 18.47 | *Artery - Aorta* | 14.63 | 9.235 | 17.31 | 58.05 |
| *Kidney - Cortex* | 6.303 | 6.632 | 6.614 | 22.18 | *Prostate* | 11.11 | 11.475 | 20.27 | 57.625 |
| *Brain - Nucleus accumbens (basal ganglia)* | 6.557 | 7.162 | 12.03 | 20.17 | *Nerve - Tibial* | 15.18 | 19.24 | 19.55 | 48.22 |
| *Heart - Atrial Appendage* | 8.609 | 5.969 | 9.644 | 22.33 | *Fallopian Tube* | 14.57 | 13.31 | 25.31 | 49.21 |
| *Brain - Cortex* | 5.5355 | 6.8145 | 11.095 | 25.555 | *Colon - Sigmoid* | 13.24 | 10 | 24.46 | 55.42 |
| *Brain - Frontal Cortex (BA9)* | 7.597 | 6.766 | 13.7 | 23.46 | *Cells - Transformed fibroblasts* | 29.98 | 34.04 | 17.95 | 25.17 |
| *Stomach* | 8.19 | 9.396 | 12.55 | 24.365 | *Brain - Cerebellum* | 9.384 | 16.21 | 12.41 | 70.97 |
| *Brain - Spinal cord (cervical c-1)* | 9.183 | 13.54 | 10.54 | 22.75 | *Spleen* | 10.935 | 25.695 | 24.555 | 49.11 |
| *Skin - Sun Exposed (Lower leg)* | 11.62 | 18.75 | 10.87 | 19.18 | *Ovary* | 12.4 | 19.46 | 20.07 | 59.51 |
| *Skin - Not Sun Exposed (Suprapubic)* | 10.72 | 18.3 | 11.7 | 19.98 | *Cervix - Ectocervix* | 14.765 | 24.625 | 23.54 | 51.435 |
| *Minor Salivary Gland* | 10.33 | 13.21 | 13.03 | 24.15 | *Brain - Cerebellar Hemisphere* | 13.325 | 17.52 | 14.98 | 68.75 |
| *Esophagus - Mucosa* | 10.58 | 24.62 | 11.92 | 18.2 | *Cervix - Endocervix* | 15.15 | 19 | 26.62 | 55 |
| *Colon - Transverse* | 9.484 | 12.52 | 14.885 | 28.96 | *Testis* | 14.12 | 49.84 | 23.42 | 31.66 |
| *Adrenal Gland* | 10.045 | 9.7865 | 12.39 | 35.965 | *Uterus* | 16.6 | 21.33 | 30.43 | 66.03 |
| *Adipose - Visceral (Omentum)* | 12.25 | 12.14 | 14.71 | 32.23 | *Cells - EBV-transformed lymphocytes* | 27.94 | 79.945 | 41.36 | 32.195 |
| *Small Intestine - Terminal Ileum* | 9.633 | 14.12 | 18.32 | 30.46 |  |  |  |  |  |

Expression of *RNASEH* genes in different 53 human tissues. Data obtained from GTEx consists of the median TPM from >90 individuals for each tissue. Blue color indicates low expression levels and red color indicates high expression levels.

**Figure 3.**
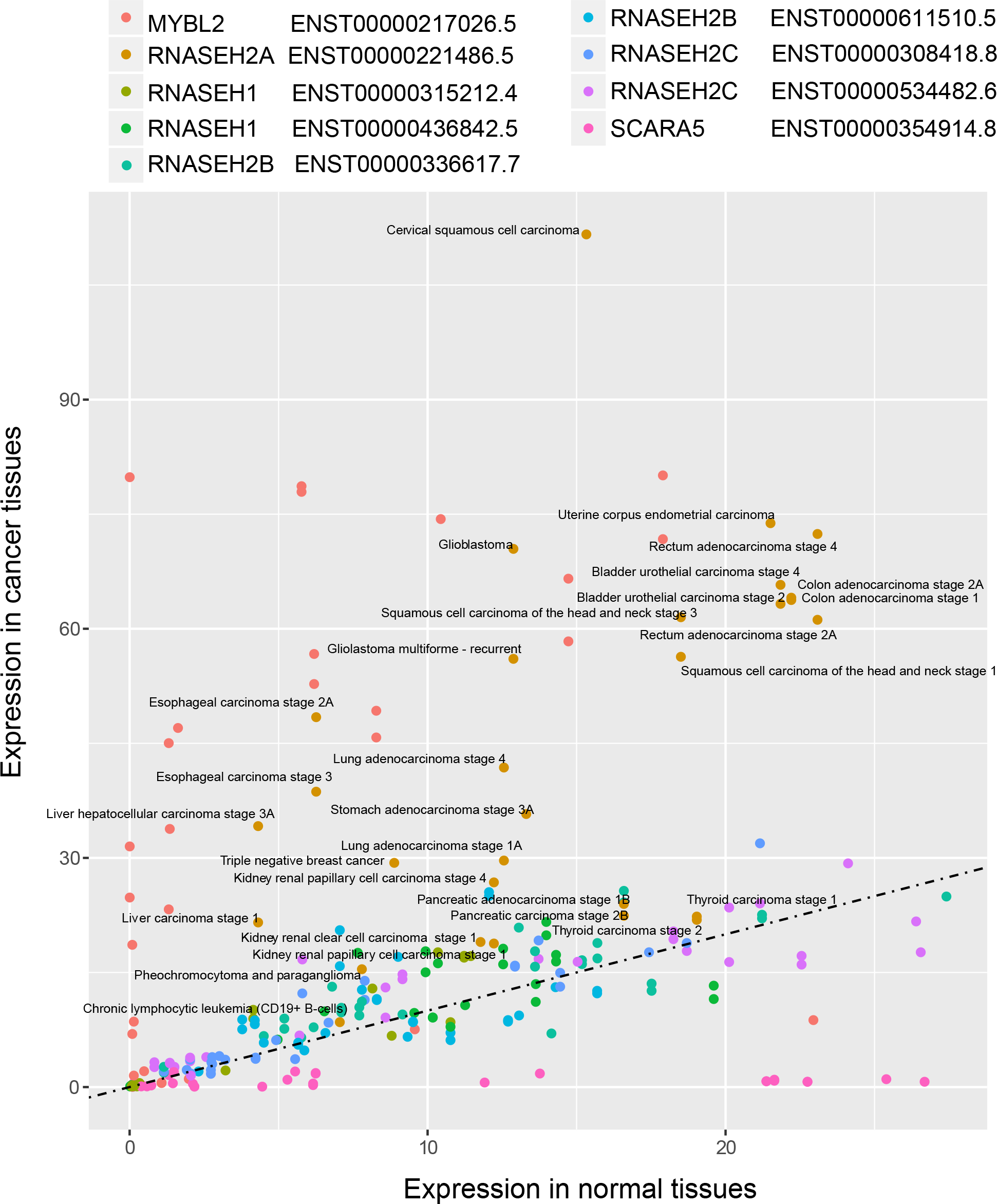
*RNASEH2A* gene expression in cancer tissues. Expression levels in TPM of different transcripts in 29 different cancer tissues compared to non-cancerous tissue controls. Data obtained from Cancer RNA-seq Nexus (CRN). Regression line represents equal expression levels in cancer and normal tissues.

**Figure 4.**
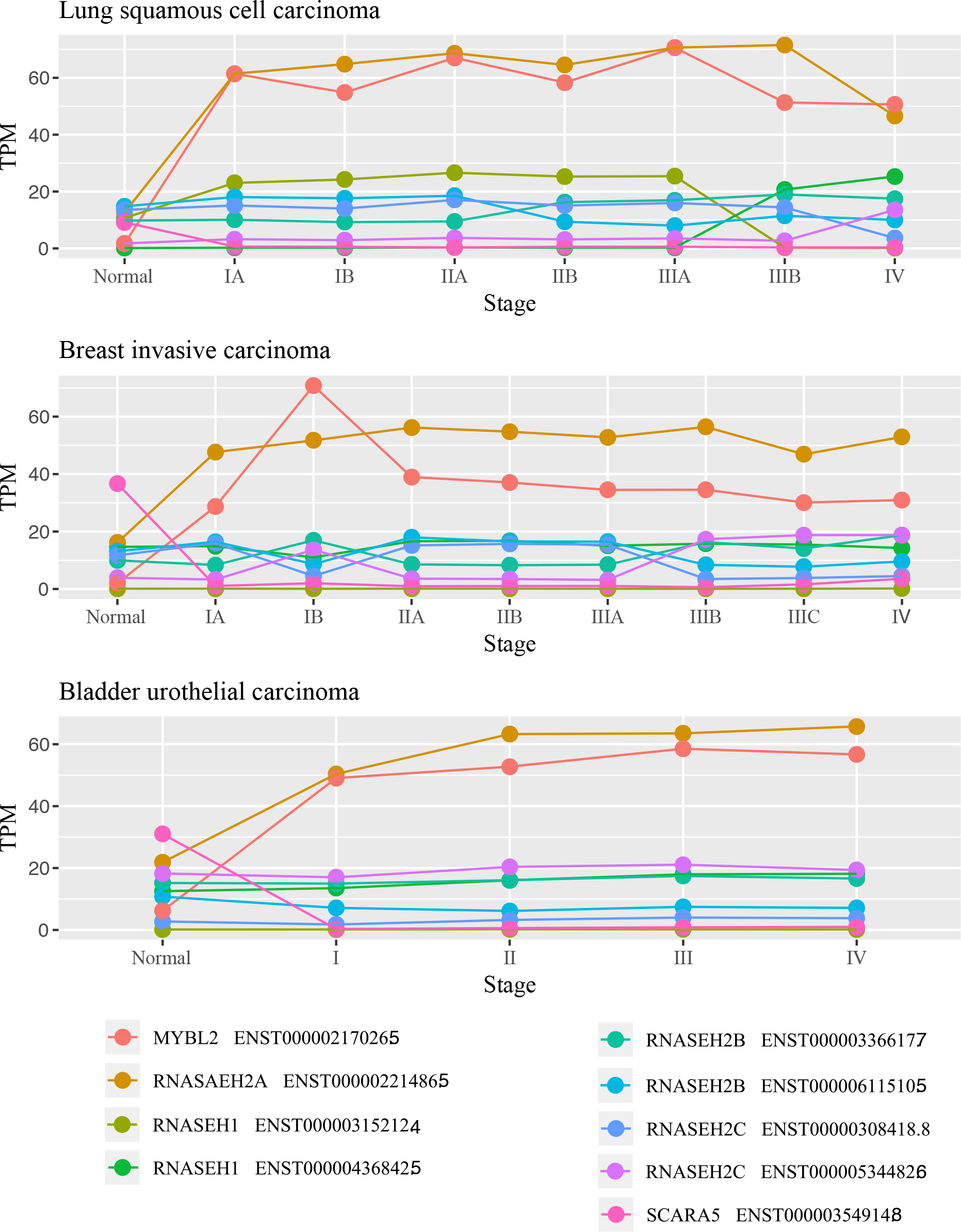
*RNASEH2A* levels are up-regulated in early stages of cancer and remain high with cancer progression. Expression levels in TPM of different transcripts of *RNASEH* genes, including *MYBL2* and *SCARA5*, was examined in normal tissues and in different stages of lung squamous adenocarcinoma, breast invasive carcinoma and bladder urothelial carcinoma (top to bottom, respectively).

## Discussion

Co-expression correlation analysis of genes is a simple approach to suggest functional network of genes. We anticipate that this approach is more accurate for analyzing genes the more their coexpressed genes share a higher correlation value. In addition, we assume that the genes analyzed should have a functional network involved in a process that is dependent on a spatio-temporal gene expression profile. Cell cycle regulation is a good example for such a process. In fact, the analysis of *CDK1*, a regulator of cyclin B implicated in cell cycle control, shows enrichment of genes involved in mitotic cell cycle regulation, microtubule cytoskeleton organization and regulation of G1/S transition of mitotic cell cycle (Supplementary Table S10 and complete list of *CDK1* coexpressed genes in Supplementary Table S11). This approach can be applied to other processes such as for example RNA regulation in stress granules, which is a well-orchestrated process dependent on time and stress conditions. In this context, *G3BP1*, an RNA helicase that is one of the key assemblers of the stress granules, shows enrichment of co-expressed genes involved in molecular function of RNA binding and RNA helicase activity in a ribonucleoprotein cellular component (Supplementary Table S10 and complete list of *G3BP1* co-expressed genes in Supplementary Table S12).

Another factor that should be considered is defining the cutoff of correlation of the co-expressed genes. In this study we arbitrary placed the cutoff on the top 2% of the co-expressed genes, in order to have a handful number of genes for analysis and to delimit false positive results. Analysis of the top 2% co-expressed correlated genes to *RNASEH1* revealed functional network of genes that are involved in RNA binding (Supplementary Table S13 for functional network analysis, and Supplementary Table S14 for complete list of *RNASEH1* co-expressed genes). After filtering top 2% correlated genes, we could identify processes such as regulation of telomerase RNA and telomeres maintenance (Supplementary Table S13), which were previously reported [32,33]. When we analyzed the top 2% co-expressed genes of *RNASEH2B*, we found the functional network of genes involved in DNA replication (Supplementary Table S15 Table for functional network analysis and Supplementary Table S16 for complete list of *RNASEH2B* co-expressed genes), as expected for this RNase H2 subunit. For *RNASEH2C* we could not find any functional network neither in 2% nor 5% analysis as there were no GO terms with an enrichment p-value below the specified value p<0.001 (data not shown). As shown in Table 3, the *RNASEH2C* expression pattern in the human tissues analyzed is different from the other *RNASEH* genes, and this perhaps explains why we could not find any functional network associated with this gene.

Remarkably, among the three *RNASEH* genes, we found that only *RNASEH2A* was involved in the functional network of mitotic cell cycle regulation. One possibility is that RNASEH2A has a function independent of RNASEH2B and RNASEH2C, as its distribution in the cell was shown not to be limited to the interaction with RNASEH2B and RNASEH2C [6]. This notion was also suggested by others showing that knocking out *RNASEH2A* in HeLa cells results in an increased sensitivity to *ATR* inhibitors compared to knocking out *RNASEH2B* [34]. We propose that *RNASEH2A* levels constitute an additional layer of regulation of *RNASEH2A* activity, which is not dictated by *RNASEH2B* or *RNASEH2C* levels, and this is why the co-expression correlation analysis did not show a functional network of mitotic cell regulation for *RNASEH2B* and *RNASEH2C* genes. Following this notion, it would be intriguing to examine the levels of the three RNase H2 genes throughout the cell cycle in human cells, and study how perturbing their levels affects RNase H2 function and/or cell cycle progression. Similar studies have been performed on yeast showing a cell cycle dependent expression of *RNH201* (orthologous gene of *RNASEH2A*), peaking at S and G2 phases [23,35], and being recruited to chromatin/telomeres at the G2 phase [35].

Rejins *et al*.[36], using an anti-mouse antibody for all proteins of the RNase H2 complex, demonstrated an increase in the expression level of RNase H2 in the mouse blastocyst, in all three embryonic layers during gastrulation and showed, in newborns and adults, that the expression becomes restricted to highly proliferative tissues such as intestinal crypt epithelium and testes [36]. Moreover, they reported that RNase H2 levels correlate with the proliferation marker Ki67 [36]. These data are consistent with our findings showing an increase in RNASEH2A protein level in actively cycling tissues as well as different cancer tissues compared to normal ones.

Using our co-expression correlation approach, we have identified three main functional networks in which *RNASEH2A* is involved: DNA replication, DNA damage repair and regulation of chromosome segregation in mitosis. Hyper-proliferation of cells results in excessive addition of RNA Okazaki fragments into the genome during DNA replication; increases the number of events when DNA polymerase mistakenly insert ribonucleotides instead of deoxy-ribonucleotides into the replicated genomic DNA [37,38]; and increases the rate of chromosomes reduction from 4N to 2N by symmetric segregation into two new daughter cells. All these events correspond to the functional networks identified in this study for *RNASEH2A* and link *RNASEH2A* activity to cell cycle regulation. In support of this notion, we have identified several cellular pathways among the highest *RNASEH2A* co-expressed genes such as the MCM complex, a replicative eukaryotic helicase that in yeast has been demonstrated to be activated upon accumulation of RNA-DNA hybrids at the G2-M checkpoint. This finding suggests that targeting the *RNASEH2A* level and/or activity could prevent the DNA damage occurring during replication which leads to mitotic catastrophe and cell death [39,40]. The other identified complexes, such as the *NDC80* complex, the *CENP* family and the *KIF* family have all been shown to function in cell cycle regulation through regulation of the microtubule filaments with the kinetochore [41,42,43].

Our mass spectrometry analysis shows 22 proteins that specifically interact with RNASEH2A. The top two proteins observed are RNASEH2B and RNASEH2C, confirming the authenticity of the results. Additional proteins that we identified in the context of RNASEH2A biology and mitosis regulation, are CCT2 and CCT8, which are part of CCT/TRiC complex. This complex was reported to regulate telomerase function by mediating its trafficking from the cytoplasm to the telomeres [26]. In addition, it was shown to promote chromosome segregation by disassembling checkpoint complex from sister chromatids in the initial phase of mitosis [27], supporting the RNASEH2A functional network in mitosis regulation.

The protein-protein interactions of RNASEH2A with nucleolin and Hsp90-beta that we detected by mass spectrometry in this study deserves further investigation. In this respect, nucleolin has been shown to interact with HSP90 in the nucleus, suggesting that HSP90 stabilizes nucleolin, which in turn, stabilizes mitotic mRNAs level linked to cancer formation [29]. As we performed mass spectrometry analysis in HEK293 cells, it would be interesting to further validate the biological significance of this interaction in normal and diseased conditions and investigate how the interaction of RNASEH2A with nucleolin and HSP90 is regulated.

HSP90, besides being recently discovered acting also in the nucleus, is mainly a chaperone protein localized in the cytoplasmic compartment [44]. Because we used a cellular system in which RNASEH2A was overexpressed, we cannot exclude that the interaction observed is the result of a different conformation and modified post-translational modification of the RNASEH2A protein, been so recognized as a “client protein” by HSP90-beta. On the other hand, the possible involvement of HSP90-beta and nucleolin together with RNASEH2A in the mitotic process could reveal important features of the biological role of RNASEH2A in the contest of genomic integrity and stability, so better defining RNASEH2A as a putative target for cancer therapy.

Our analyses of expression levels focused on the *RNASEH* genes (*RNASEH1, RNASEH2A, RNASEH2B* and *RNASEH2C*) using the GTEx v.7 database revealed a marked contrast in *RNASEH2A* expression levels between low and high proliferative tissues. *RNASEH2A* showed low expression levels in low proliferative tissues such as brain, kidney (cortex), heart and whole blood, and high expression levels in highly proliferative tissues, such as skin, esophagus, small intestine, testis, cervix, as well as in transformed fibroblasts and transformed lymphocytes (Table 3). Moreover, in line with the initial study by Flanagan et al. [18], showing elevated expression levels of *RNASEH2A* in transformed mesenchymal stem cells, our analysis of 29 different randomly selected tumors in different tissues compared to their healthy controls, revealed that *RNASEH2A* is the only *RNASEH* gene displaying increased expression levels (Figure 3). Further analyses of *RNASEH* expression levels throughout different stages of lung squamous adenocarcinoma, breast invasive carcinoma and bladder urothelial carcinoma, again highlighted specific upregulation of *RNASEH2A* from early stages to more advanced stages of these cancers (Figure 4). It is interesting to note that Dai et al., reported on a possible role of *RNASEH2A* in glioma cell proliferation. Their findings suggested a role of *RNASEH2A* upregulation in cell growth and apoptosis, contributing to gliomagenesis and cancer progression [45]. More recently, *RNASEH2A* overexpression was associated with cancer cell resistance to chemotherapy *in vitro*, and with aggressiveness and poor outcomes in breast cancers of ER-positive subtypes [46].

Overall, our study highlights an emerging role of RNASEH2A in cell cycle regulation, that appears independent from its function as part of the RNase H2 whole enzyme. The presented findings stimulate new research directions to better understanding and characterizing the function of RNASEH2A in cell proliferation both in healthy and in cancer cells, and support further exploring RNASEH2A as target for cancer diagnosis and therapy.

## Materials and methods

### Cell and protein extracts

HEK293 were grown in DMEM media (Corning, Corning NY, cat# 45000-304), containing 10% FBS (Sigma-Aldrich, St. Louis, MO, cat # F0926) and 1% Penicillin-Streptomycin (Fisher Scientific, Hampton NH, cat # 15-140-122) in 5% CO_2_ incubator at 37 °C and were passaged once a week to maintain cell growth. Whole cell lysate from HEK293 was extracted with NP-40 lysis buffer (50mM Tris-HCL pH 7.4, 150mM NaCl, 0.5mM EDTA, 0.5% NP-40) together with a cocktail of protease inhibitor (Complete EDTA-free, Roche Applied Science, Indianapolis, IN) followed by centrifugation at 14,000 rpm for 15 minutes at +4 °C to pellet the debris.

### Plasmids and transfection

To over-express *RNASEH2A* in HEK293 cell line we used the pEGFP-RNASEH2A plasmid. (Addgene, Watertown MA, cat # 108700). As control, we cloned the eGFP plasmid by restricting the pEGFP-RNASEH2A using HincII (New England Biolabs, Radnor, PA, cat # R0103S) that cut upstream and downstream of the RNASEH2A gene but leaves the EGFP intact. Then, the plasmid was ligated by T4 ligase (New England Biolabs, cat# 101228-176) and transformed into Agilent XL1-Blue Electroporation-Competent Cells (Agilent, Santa Clare, CA, cat # 50-125-045) by electroporation using Gene Pulser Xcell electroporation system (Bio-Rad, Hercules, CA, cat# 1652660). Isolated colonies were grown in Luria-Bertani (LB) medium over-night; plasmids were extracted using the miniprep kit (Fisher Scientific, cat # FERK0503) and sent to DNA sequencing to verify the plasmid sequence.

For transient transfection, HEK293 cells were plated at 1-3×10^6^ per 10cm plate. The following day 5μg of the plasmids were incubated with 25μl of the Lipofectamine transfection reagent (Fisher Scientific, cat # 11668019) for 10-15 minutes at room temperature in Opti-MEM media (Fisher Scientific, cat # 31985062) and the plasmid-reagent mix was distributed to the cells. The next day transfected cells were fed with fresh media and harvested 48 hours after transfection.

### Co-immunoprecipitation

Whole cell lysate (1mg) from HEK293 cells transfected with either the eGFP or the pEGFP-RNASEH2A plasmid was incubated with 30 ul of GFP-Trap MA coupled to magnetic particles (ChromoTek, Planegg-Martinsried Germany, gtma-10 lot # 80912001MA) at 4 °C with end to end rotation for 1 hour. Then, the magnetic beads-GFP protein complex were separated from the rest of the protein extract using a magnetic stand. The beads were washed twice with wash buffer I (50mM Tris-HCL pH 7.5, 150mM NaCl, 0.25% NP-40) and additional 2 washes with wash buffer II (50mM Tris-HCL pH 7.5, 150mM NaCl). Then, the beads were resuspended in SDS loading buffer and boiled at 95 °C for 10 minutes. Finally, the beads were centrifuged at 13000 RPM for 5 minutes at room temperature and the eluted proteins were collected and used for Western blot or mass spectrometry analysis.

### Protein analysis (Western blot)

Total protein concentration from HEK293 lysate were quantified (Bradford Protein Assay, Pierce Biotechnology, Rockford, IL) and 15ug of the protein extracts were separated on a 10% SDS–polyacrylamide electrophoresis gel, transferred to 0.45μm nitrocellulose membrane (Amersham, Buckinghamshire, UK cat #10120-006). The blotted membrane was then blocked in 5% non-fat dry milk TBS-T (10mM Tris-Cl [pH 7.5], 100mM NaCl, 0.1% Tween-20) at room temperature for 1 hour and incubated over-night with the specific primary antibody: anti-RNASEH2A (Santa Cruz Biotechnology, Dallas TX, cat # SC-515475, 1:1000 lot # B2818), anti-GFP (B2) (Santa Cruz Biotechnology, Dallas TX, cat # SC-9996, 1:1000 lot # C0619. After washing 3x with TBS-T, the membrane was incubated with a mouse secondary antibody conjugated with horseradish peroxidase (Thermo Fisher Scientific, Waltham, MA, cat #31430 1:10000at room temperature for 1 hour. Following washing with TBS-T, protein signals were visualized using the ECL method according to the manufacture’s recommendations (PierceTM ECL Western Blotting Substrate, Thermo Fisher Scientific cat# 32109) and exposed on autoradiograph films (Denville Scientific, South Plainfield, New Jersey cat# 3018).

### Mass spectrometry analysis

#### Sample Preparation, Trypsin Digestion, LC-MS/MS, and Peptide Identification

The gel lanes were excised from the gel and the proteins were reduced, alkylated and digested with trypsin as previously described [47] with the following modification. Each gel lane was only divided into four sections and then pooled following digestion. The peptides were analyzed by nano-LC-MS/MS, and peptide identification as previously described [48]. The Raw files were searched using the Mascot algorithm (ver. 2.5.1) against a protein database constructed by combining the bovine reference database (Uniprot.com, 37,941 entries, downloaded 5-22-19), the human reference database (Uniprot.com, 73,911 entries, downloaded 5-22-19), the GFP-RNASEH2A protein, and a contaminant database (cRAP, downloaded 11-21-16 from http://www.thegpm.org) via Proteome Discoverer 2.1. A 1% FDR (“High Confidence”) was used for both the peptides and the proteins. At least of 3 peptide sequences for the RNASEH2A protein were considered for the analysis.

### Co-Expression Correlation analysis

Data obtained from GTEx portal v7 [3] (https://gtexportal.org/home/datasets) was the RNA-seq data consisted the gene median TPM (transcripts per million) by tissue. The data contained the expression levels of 56,202 transcripts. All the transcripts that were not transcribed in at least one tissue were eliminated, and there were 42,548 transcripts remaining. Then, we used the excel formula (=correl(dataset(RNASEH2A), dataset(individual gene in the database))) to correlate RNASEH2A expression with all 42,547 transcripts among all 53 tissues. Then, we analyzed the GO-term of the top 2% co-expressed genes (851 genes) using Gene Ontology enRIchment anaLysis and visuaLizAtion tool (GOrilla) [24].

### Co-expression correlation analysis verification with STRING database

The top 9 gene with highest correlation with *RNASEH2A* (including itself) were analyzed with the STRING protein network association database V11.0 [25]. We decided to use only genes/proteins for which we could identify 20 other proteins associated in the same network in the organism *Homo sapiens* using the settings Textmining/Experiments/Databases/Gene Fusion and excluding Co-expression/Neighborhood/Co-occurrence. We examined proteins interactions on the 1^st^ shell that refers to proteins directly associated with the input protein, using a medium confidence score of 0.400. The 139 identified proteins were aligned with the co-expression correlation list for RNASEH2A.

### Gene Expression analysis in cancer

Data obtained from Cancer RNA-Seq Nexus (CRN), database providing phenotype-specific coding-transcript/lncRNA expression profiles and mRNA–lncRNA co-expression networks in cancer cells (30) were used to examine the expression level of *RNASEH* genes in cancer.

### Data availability

The raw data of the study are presented within the Supplementary Tables and are also available from the corresponding author.

## Acknowledgments

We thank Georgia Institute of Technology’s Parker H. Petit Institute for Bioengineering and Biosciences including the Systems Mass Spectrometry Core Facility for supporting this work, C. Meers and D. L. Kundnani for critical reading of the manuscript, and all the people of the Storici laboratory for assistance and feedback on this research. We acknowledge funding from the National Institutes of Health, NIH, NIGMS R01 GM115927 (F.S.); the National Science Foundation, NSF, MCB-1615335 and the Howard Hughes Medical Institute Faculty Scholar grant 55108574 (F.S.).

## Author contributions

A.T. and F.S. conceive and design experiments. A.T. conducted the experiments, performed most of the data analyses, and wrote the first draft of the manuscript. S.M. assisted and revised data analysis and completed the preparation of the manuscript with support by F.S.

## Additional Information

### Competing interests

The authors declare no competing interests.

## Supplementary information

### Table legends

**Supplementary Table S1.** List of *RNASEH2A* co-expressed correlated genes.

**Supplementary Table S2.** Gene ontology of the 2% co-expressed correlated *RNASEH2A* genes using the GOrilla analysis tool.

**Supplementary Table S3.** Protein network association data obtained from STRING proteinprotein association network applied to RNASEH2A and its top co-expressed correlated genes.

**Supplementary Table S4.** List of the protein interactors for each of the top 9 *RNASEH2A* coexpressed genes aligned with the co-expressed *RNASEH2A* genes.

**Supplementary Table S5.** List of proteins identified by mass spectrometry.

**Supplementary Table S6.** Expression level of the *RNASEH* genes in cancer using data obtained from Cancer RNA-seq Nexus (CNR) in 30 different randomly selected tumors.

**Supplementary Table S7.** Expression level of the *RNASEH* genes in cancer using data obtained from Cancer RNA-seq Nexus (CNR) in different tissues at different stage of cancer progression.

**Supplementary Table S8.** Level of the *RNASEH* genes throughout the progression of cancer showing constant upregulation of *RNASEH2A* in lung, breast and bladder tumors.

**Supplementary Table S9.** Level of the *RNASEH* genes at different stages of cancer.

**Supplementary Table S10.** Gene ontology of the 2% co-expressed correlated *CDK1* genes and *G3BP1* using the GOrilla analysis tool showing enrichment of genes involved in cell cycle regulation and in molecular binding of RNA.

**Supplementary Table S11.** List of *CDK1* co-expressed correlated genes.

**Supplementary Table S12.** *G3BP1* co-expressed correlated genes.

**Supplementary Table S13.** Gene ontology of the 2% and 5% co-expressed correlated *RNASEH1* genes using the GOrilla analysis tool showing a functional network of genes involved in RNA binding and processes of regulation of telomerase RNA and telomeres maintenance.

**Supplementary Table S14.** List of *RNASEH1* co-expressed correlated genes.

**Supplementary Table S15.** Gene ontology of the 2% co-expressed correlated *RNASEH2B* using the GOrilla analysis tool showing functional network of genes involved in DNA replication. **Supplementary Table S16.** List of *RNASEH2B* co-expressed correlated genes.

## Notes

### Competing Interest Statement

The authors have declared no competing interest.

